# Autoinducer-2 and acyl homoserine lactones have contrasting effects on ammonia and nitrite-oxidizing sludge

**DOI:** 10.64898/2026.01.08.698431

**Authors:** Hira Waheed, Yuchen Zhang, Lan Nguyen, Abigail S. Joyce, Lee Ferguson, Jeseth Delgado Vela

## Abstract

Enhancing nitrification with quorum sensing manipulation has emerged as a promising strategy to overcome rate-limiting steps. This study examined how disrupting microbial cell-to-cell communication by supplementing or quenching signal molecules regulates ammonia– and nitrite-oxidizing activity within activated sludge. Prolonged enrichment of activated sludge over 180 days yielded stable and robust nitrifying consortia, increasing the ammonia oxidation rate (AOR) from 5.4 to 9.3 mg N g⁻¹ VSS h⁻¹ and the nitrite oxidation rate (NOR) from 0.6 to 4.8 mg N g⁻¹ VSS h⁻¹, while reducing the hydraulic residence time by 50 % (from 72 h to 36 h). Exogenous addition of oxoacyl and long-chain acyl homoserine lactone (AHL) signals further boosted AOR up to 4.5-fold higher than the control activated sludge, predominantly through transcriptional activation of the *amoA* gene in *Nitrosomonas eutropha*. Acylase-mediated AHL quenching lowered AOR to 4.8 mg N g⁻¹ VSS h⁻¹ but improved functional resilience of nitrite oxidizing bacteria by enhancing mass transfer and oxygen diffusion via smaller flocs (123 µm vs 357 µm in AHL-treated sludge). Conversely, elevated autoinducer-2 (AI-2) levels suppressed ammonia-oxidizing activity (AOR = 2 mg N g⁻¹ VSS h⁻¹) yet stimulated nitrite oxidation (NOR = 36 mg N g⁻¹ VSS h⁻¹), particularly *Nitrospira*, underscoring the contrasting regulatory requirements of the two nitrifying guilds. Overall, the study demonstrates that AHLs and AI-2 serve as complementary yet opposing regulators of nitrification, primarily activating ammonia oxidizers and nitrite oxidizers, respectively. Maintaining these signaling molecules within an optimal physiological window thereby offers a biologically tunable approach for synchronized ammonia and nitrite oxidation, ultimately maximizing nitrogen removal in biological treatment systems.

## 1. Introduction

Nitrogen removal via nitrification and denitrification is a widely recognized conventional process for wastewater treatment systems to address eutrophication. Nitrification, a primary step in the nitrogen cycle, is predominantly regulated by autotrophic nitrifiers. This involves ammonia-oxidizing archaea (AOA) and ammonia-oxidizing bacteria (AOB) mediating conversion of ammonia to nitrite, followed by the oxidation of nitrite to nitrate via nitrite oxidizing bacteria (NOB). Comammox bacteria, which convert ammonia to nitrate, can also be involved in nitrification processes. Although the overall efficiency of nitrification in wastewater treatment systems is strongly influenced by both biological and operational factors, the proliferation of nitrifying microorganisms is a critical rate-limiting step towards nitrogen removal. Nitrifiers have slow growth kinetics, which hinders rapid startup-up and leads to prolonged solids/hydraulic retention times, increasing energy requirements (Feng et al., 2019). Sustainably enhancing the activity of nitrifiers to ensure treatment stability is crucial to ensure the reliability of nitrification-based treatment systems.

Quorum sensing (QS), i.e., cell-to-cell microbial communication has gained attention as a mechanism that can be used to augment nitrifying activity. QS allows bacteria to synchronize responses across a population and modulate social behaviors via chemical signals (autoinducers). Reported autoinducer classes include, but are not limited to, acyl-homoserine lactones (AHLs), autoinducer-2 (AI-2), autoinducer-3, and autoinducer peptides (Lu et al., 2022). Since nitrifying bacteria are predominantly gram-negative, recent investigations have focused on AHL– and AI-2–mediated QS mechanisms as potential means to enhance nitrification (Chen et al., 2023; Shourjeh et al., 2021). Feng et al. (2019) correlated successful nitritation with endogenously produced AHLs in activated sludge. Exogenous supplementation of two AHLs (C4-, and C6-HSL), in combination with free ammonia, enhanced start-up efficiency, and maintained stable partial nitrification with 44% increase in AOB abundance (Zhao et al., 2025). In an integrated floating fixed-film activated sludge system, the influence of AHL mixtures (C4-, C8-, 3OHC12-, and C14) on simultaneous nitrification and denitrification (SND) was examined, revealing higher biomass density with enhanced nitrifying and denitrifying bacterial activity (Liu et al., 2021). Similarly, increase in nitrite accumulation via AI-2 was observed during partial nitrification (Hu et al., 2022). The autoinducers improved nitrifying activity by stimulating energy metabolism, promoting microbial communication, increasing extracellular polymeric substance production for structural stability, and amplifying QS responses by triggering endogenous signal production in response to exogenous signals (Gao et al., 2025; Liu et al., 2021; Zhao et al., 2025).

Contrary to QS, quorum quenching (QQ) suppresses microbial communication by inactivating autoinducers (Chen et al., 2023). Various enzymes, including lactonases, acylases, kinases, and oxidoreductases as well as chemical compounds such as vanillin, quercetin, flavonoids and furanones, have been reported to interfere with QS pathways, either as metabolites naturally produced by microorganisms or supplemented externally (Chen et al., 2023; Waheed et al., 2021). In wastewater treatment plants, QS and QQ processes inevitably coexist. Therefore, understanding the balance between QS and QQ is critical, as QS can stimulate nitrifying activity and structural stability, whereas QQ may suppress excessive signaling and regulate community composition, collectively shaping the efficiency and resilience of nitrification processes.

The existing literature on QS-based nitrification has primarily focused on community composition via metagenomic approaches, while largely overlooking the link between autoinducer exposure and functional gene markers of dominant AOB and NOB populations that could reveal the specific mechanism connecting QS to nitrogen-metabolism pathways. Most studies have tested only a limited number of AHLs or focused exclusively on either AHLs or AI-2, neglecting the broader spectrum of signaling molecules in wastewater. In addition, earlier work has generally evaluated the influence of autoinducers on nitrifying sludge as a whole, without distinguishing the individual responses of AOB and NOB. However, separate testing of these sludges is needed to clearly assess autoinducer driven shifts and to better understand functional balance of each community within mixed nitrifying systems. Likewise, studies on QQ interventions during nitrification remain limited. Altogether, these gaps highlight the need for a comprehensive evaluation encompassing a broad range of autoinducers and their respective quenching effects to understand their collective influence on ammonia and nitrite oxidation processes, as well as their regulatory control over key nitrifying functional genes.

The present study aimed to elucidate the intrinsic relationship between QS, QQ, and nitrogen metabolism by employing a broad spectrum of autoinducers and simultaneously tracking temporal dynamics of key nitrifiers in terms of gene abundance and transcriptional activity of functional genes. We systematically examined ammonium and nitrite transformation in AOB– and NOB-enriched sequencing batch reactors with (a) disruption of microbial coordination by quenching endogenous autoinducers and (b) enhancement of cell-to-cell communication by supplementing nine sludge-derived AHLs and an AI-2 molecule. The findings highlight the potential of autoinducer-based manipulations as promising biological strategies to achieve rapid start-up and stable performance at lower hydraulic residence time (HRT), thereby advancing the development of sustainable and robust full-scale nitrogen removal technologies.

## 2. Materials & Methods

### 2.1. Chemicals and reagents

All AHLs used in this study were purchased from Sigma-Aldrich (USA), including N-butyryl-(C4-), N-hexanoyl-(C6-), N-3-oxohexadecanoyl-(3OC6-), N-heptanoyl-L-(C7-), N-octanoyl-(C8-), N-3-oxooctanoyl-(3OC8-), N-decanoyl-(C10-), N-3-oxodecanoyl-(3OC10-), N-dodecanoyl-(C12-), and N-oxododecanoyl-(C12-) HSL. In addition, N-butyryl-L-HSL-d5 (C4-HSL-d5), and N-hexanoyl-L-HSL-d3 (C6-HSL-d3) from Cayman Chemicals (USA) were used as internal standards for the study. A linear precursor of AI-2 4,5-Dihydroxypentane-2,3-dione (DPD), its derivatizing reagent dimethyl 2,2ߣ-(4,5-diamino-1,2-phenylene)bis(oxy)diacetate, and [4-^13^C]-DPD as an internal standard were procured from Rita Ventura’s research group at Instituto de Tecnologia Química e Biológica (Oeiras, Portugal). Acylase I from porcine kidney and Quercetin purchased from Sigma-Aldrich (USA) were used as quenchers in the study.

### 2.2. Bioreactors operation

The activated sludge used to seed the enrichment reactors was obtained from a water reclamation facility in North Carolina with a four-stage Bardenpho process with an initial mixed liquor suspended solids (MLSS) concentration of ∼6000 mg L^−1^. Four independent sequencing batch reactors, each with an effective volume of 1.8 L, were operated to acclimate the seed sludge for over a year with a 12-h cycle, consisting of 30 min feeding, 10 h mixing and aeration, 1 h settling, and 30 min for water discharge. Two sets of duplicate bioreactors were designated for separate cultivation of AOB and NOB, with influent nitrogen maintained at 200 mg L^−1^ as 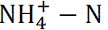 or 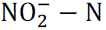 by supplementing (NH_4_)_2_SO_4_ or NaNO_2_, respectively. Other than the nitrogen source, the remaining composition of the synthetic feed was identical across all bioreactors and is listed in Table S1. The pH was maintained for AOB-bioreactors to 7.5±0.3 via continuous dosing of NaHCO_3_ throughout the cultivation period. All bioreactors were operated at hydraulic retention time and solids retention time of 10 h, and 35 days, respectively with a DO level of 4.3 ± 0.5 mg O_2_ L^−1^.

### 2.3. Experimental design

All batch experiments were performed in duplicates using sterile conical tubes (50 mL), at room temperature with an orbital shaker at 100 rpm. Prior to each batch test, nitrifying sludge was washed three times with deionized water to remove residual substrates. The concentrations of MLSS and mixed liquid volatile suspended solids (MLVSS) during batch tests were maintained to 500 ± 50 mg L^−1^ and 400 ± 30 mg L^−1^, respectively.

#### 2.3.1. Batch test for the enrichment of AOB & NOB

Sludge from bioreactors was categorized into three stages based on adaptation/cultivation period: inoculum (day 1), transitional (day 60-65), and mature stage (day 160-180) for nitrifying community. During each stage, sludge samples (40 mL) were collected from each bioreactor in duplicate and washed three times with deionized water to eliminate residual nitrogenous substrates. Synthetic feed was added to each sludge fraction with N concentration to 200 mg-N L^−1^ using (NH_4_)_2_SO_4_ for AOB and NaNO_2_, for NOB testing. Under continuous shaking at 80 rpm, sludge samples were collected at various time points from 0 h to 96 h and centrifuged for 15 min at 4000 rpm and 4 °C. Collected supernatants and pellets were stored at −20 °C and −80 °C until further testing for nitrogen species 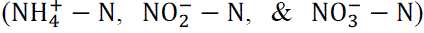 and biological analyses respectively. Of note, NaHCO_3_ was used to adjust the pH to approximately 7.5 following each sampling point for all AOB experiments. During NOB experiments the pH was at 7.98±0.24 without the need for adjustment.

After nitrifying adaptation, the sludge was considered suitable for evaluating the effects of autoinducers on AOB and NOB activity. In total, one AI-2 and nine AHLs were categorized based on their abundance in nitrifying enriched sludge and structural diversity into various sets as detailed in following sections.

#### 2.3.2. Batch test for the effects of AHLs mediated QS and QQ

Treatment sets (1-11) for AHLs were conducted separately for AOB and NOB enriched sludge. These include no addition batches (1) control with fresh sludge (CnF), and (2) control with enriched sludge (CnEn), and batches dosed with AHL of varied abundances (determined in 2.3.1) including (3) low abundant AHLs (Low), (4) moderately abundant AHLs (Medium), (5) dominant AHLs (High). Subsequent batch experiments also tested (6) a mixture of nine AHLs (All), (7) short-chain AHLs (Short chain), (8) long chain AHLs (Long chain), (9) AHLs with oxo functional group (Oxoacyl), (10) AHLs without any functional group (Acyl), and (11) Acylase as an AHL quencher (Acylase). Notably, dosing AHLs mixtures prepared for each set were first added to empty conical tube, after which the acetonitrile solvent was completely evaporated via nitrogen blowing to eliminate potential inhibitory effects on nitrification (Yu et al., 2018). The final concentration of each autoinducer in each batch was 500 nmol/L (1 nmol mg^−1^VSS), and 0.1 mg/mL of acylase in set 10 (Xiong et al., 2024). Sludge samples were collected at various time points, centrifuged, and stored as indicated in Section 2.3.1. The pH was controlled to 7.5±0.3 to avoid AHLs degradation, considering that the stability of AHLs is compromised under alkaline conditions (Chen et al., 2023), while acidic environment inhibits nitrification.

#### 2.3.3. Batch test for the effect of AI-2 mediated QS and QQ

For AOB and NOB sludge, five experimental sets were designed and conducted as (1) control with fresh sludge (CnF), (2) control with enriched sludge (CnEn), (3) DPD precursor of AI-2 (AI-2), (4) quercetin (Qc), and (5) quercetin and acylase (Qc+Acylase). Quercetin is a flavonoid polyphenol, a widely used AI-2 inhibitor (Li et al., 2025). Of note, set 5 was conducted to quantify the combined effects of quenchers targeting both AI-2 and AHLs mediated QS. The final concentration of AI-2 and each quencher was kept as 500 nmol/L, and 0.1 mg/mL, respectively.

### 2.4. Extraction and quantification of autoinducers

AHLs were extracted from the nitrifying sludge via liquid-liquid extraction as per the method described previously with slight modifications (Waheed et al., 2017). Samples were analyzed by ultra-high pressure liquid chromatography-high resolution mass spectrometry (UHPLC-HRMS) (UHPLC: Vanquish; HRMS: Fusion Lumos; both from ThermoFisher Scientific). The extraction and detection of AHLs are detailed in SI-1.2.

The AI-2 method was based on the measurement of DPD with UHPLC-HRMS following the method of (Xiao et al., 2019). Briefly, supernatant samples were centrifuged and filtered through a 0.45-µm membrane filter. 50 μL of the sample was added to an HPLC vial containing 2.5 μL of 1 mg/L ^13^C-DPD as the internal standard, followed by 5 μL of 4 g/L of tagging reagent to the vial to derivatize DPD into a quinoxaline, dimethyl 2,2’-(2-(1,2-dihydroxyethyl)-3-methylquinoxaline-6,7-diyl)bis(oxy)diacetate. The reaction was allowed to continue at room temperature for an hour and immediately analyzed via UHPLC-HRMS. The detection of AI-2 is detailed in SI-1.3.

### 2.5. Nucleic acid extractions and molecular analysis

The RNA and DNA from sludge samples were extracted using RNeasy PowerSoil total RNA and DNA elution kit (Qiagen), respectively as per manufacturer’s instructions with slight modifications (vortex time in power bead adapter was reduced to 4 min on a MP Biomedicals Fast-Prep 24 bead beater). DNA and RNA were eluted in 100 µL of elution buffer. The RNA eluted samples were further processed for DNA removal using DNA-free kit (Invitrogen, Thermo Fisher Scientific) to ensure no DNA content in the RNA samples. Nanodrop^TM^ One (Thermo scientific^TM^) and Qubit fluorometer (Invitrogen, Thermo Fisher Scientific) were used to quantify and quality check the eluted DNA and RNA concentration in the samples before ddPCR/qPCR analysis.

#### 2.5.1. Droplet digital PCR assay for functional genes

The sets of primers and probes used in the study are detailed in Table S3. Biological duplicates from the batch studies at 4 h and 12 h were selected for absolute quantification of targeted organisms, including *amoA* functional genes of *Nitrosospira* sp*., Nitrosomonas eutropha*, and *N. europaea* as AOB community, whereas 16S for *Nitrobacter* and *Nitrospira* sp. were quantified as dominant NOB. DNA and RNA extracts were diluted 1:10^4^ and 1:250, respectively, in nuclease-free water prior to quantification. Details of the reaction mixture preparation and thermal cycling conditions are provided in supplementary information (SI-1.4, Table S4).

The measurements from technical replicates were averaged and the final concentrations were converted from copies/µL to copies/mg-biomass.

#### 2.5.2. Quantitative PCR assay for housekeeping genes

Gene abundance and expression levels were normalized against housekeeping genes to account for the variations in total community. The qPCR analyses were carried out in two sets based on target community, wherein standards for *rpoB* and 16S rRNA gene primer sets were prepared using highly purified DNA extracts from AOB and NOB enriched samples, respectively, as described in text SI-1.5.

Standards for qPCR were prepared by performing a 10-fold serial dilution of the amplified target DNA ranging from 10^9^ to 10^2^ copies/µL. Each reaction was performed in triplicates using a 20 µL reaction volume containing EvaGreen Mastermix (10 µL of 2x), primer mix, nuclease-free water, and corresponding DNA standard/sample. The qPCR thermal cycling conditions are detailed in Table S5. All samples were tested in technical triplicates. Standard curves were generated for each qPCR run and only those with the amplification efficiency >85% were considered valid for quantitative analysis.

### 2.6. Analytical methods

The MLSS, and volatile suspended solids (VSS) were analyzed as per the Standard Methods (APHA, 2012). The concentration of 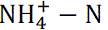 was determined using phenate method and quantified via microplate reader (APHA, 2012). Ion chromatography (Metrohm 930) was used for the quantification of 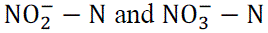 using NaNO_2_ and KNO_3_ as standards, respectively.DO and pH were monitored with a multimeter (Mettler Toledo). The particle size distribution of the sludge was measured using a laser particle size analyzer (Malvern). All tests were performed in triplicate.

### 2.7. Statistical analysis

ANOVA with subsequent Tukey comparison tests, t-tests, and Pearson Correlation Coefficient (PCC) were performed for multiple comparisons using OriginPro (Academic, 2025). Differences were considered statistically significant at *p* < 0.05.

## 3. Results and Discussion

### 3.1. Enhanced nitrification was achieved through enrichments of indigenous nitrifying communities

Activated sludge with MLSS and MLVSS concentrations of 5800±420 mg/L and 4400±320 mg/L, respectively demonstrated an ammonium removal efficiency (ARE) in the range of 28 to 55% when supplied with 200 mg/L of 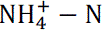 as the sole nitrogen source in a synthetic feed. While a portion of the sludge exhibited nitrite removal of 38% when fed with 200 mg-N/L of 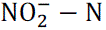 as the only nitrogen source. Given the lower growth rates of nitrifiers relative to heterotrophs, increasing their proportion within the sludge was prioritized before initiating subsequent experiments (Feng et al., 2019). Based on ammonium– and nitrite-oxidizing capacities of the initial inoculum, the activated sludge was seeded into four bioreactors, each pair dedicated to cultivating either AOB or NOB, hereafter referred to as AOB-bioreactor and NOB-bioreactor, respectively. The overall cultivation period was divided into three distinct stages: initial inoculum (I), transitional (II), and mature (III) stage for nitrifying community.

#### 3.1.1. AOB enrichment resulted in a mixture of *Nitrosomonas europaea* and *Nitrosomonas eutropha,* with *N. eutropha* displaying higher activities

The sludge consortium with an average MLSS of 500 mg/L from each stage was tested for ammonia oxidation potential when spiked with 200 mg-N/L. The initial inoculum during stage I had an ARE of 55% and increased to 82% and 100% in 36 h during stages II and III, respectively (Fig. S1a). During 180-d cultivation period, a significant difference in ammonium oxidation rate (AOR) (*p*<0.05) from stage I to III (5.4±0.25 to 9.24±0.5 mg g^−1^VSS h^−1^) was achieved (Fig. 1). This improvement likely resulted from the extended cultivation time, which is crucial for the slow-growing autotrophic nitrifiers, enabling increased nitrogen removal with reduction in overall HRT (Huang and Cui, 2025). A substantial decrease in HRT, from 72 h to 36 h, was observed from initial inoculum to well adapted consortium for complete ammonium oxidation in AOB-bioreactor.

**Fig. 1.**
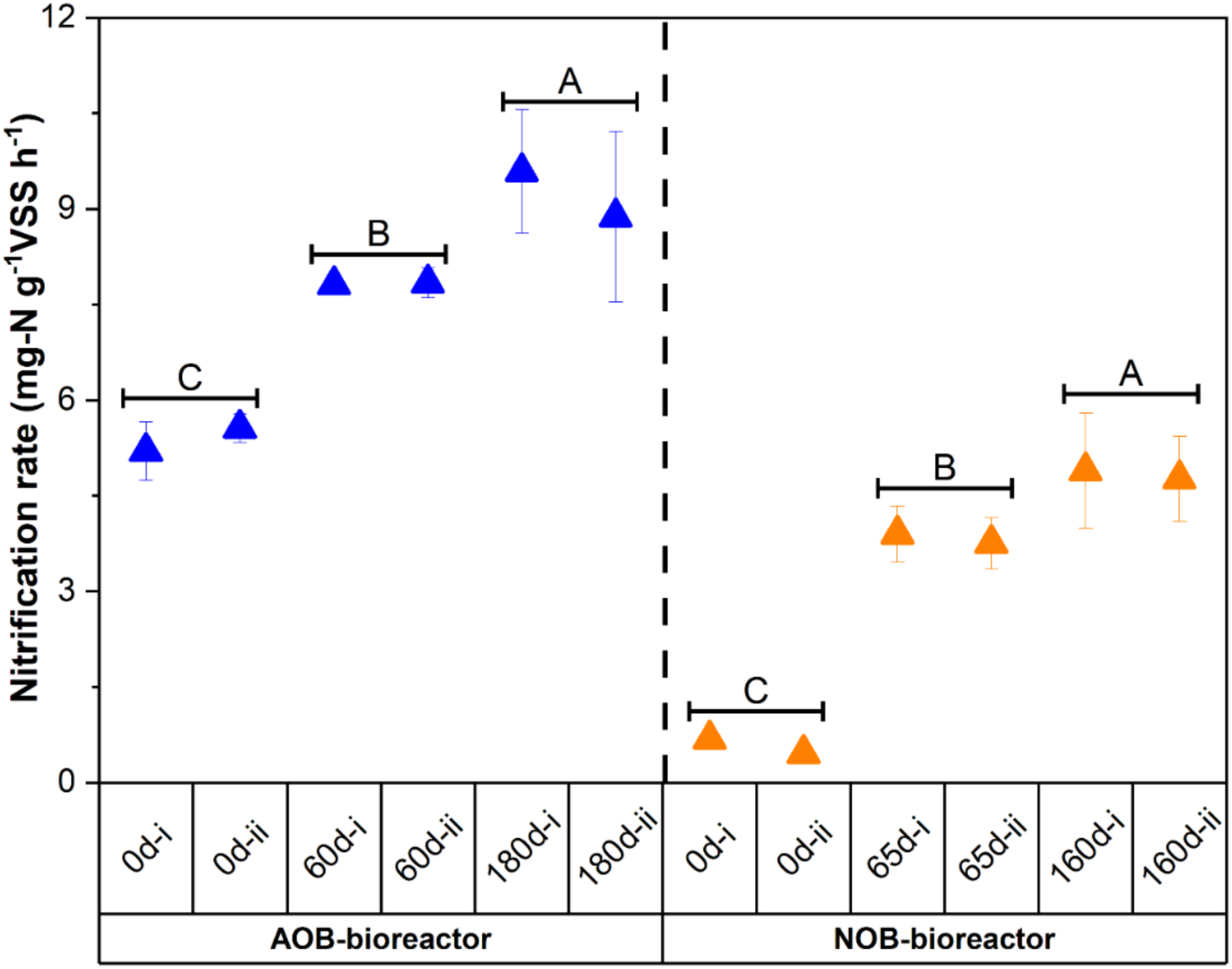
Nitrification rate measured by a linear fit of oxidized nitrogen indicating 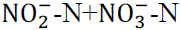 for AOB (blue), and 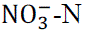 for NOB (orange). Biological replicates are grouped by horizontal brackets, and groups not sharing the same letter (A, B, C) differ significantly as per Tukey’s test (*p*<0.05).

For each cultivation stage, an *amoA* encoding ammonia monooxygenase was selected as a key functional marker linking the abundance and transcriptional activity of AOB to ammonia oxidation potential (Johnston et al., 2023). Despite targeting *Nitrosospira* sp., this lineage was absent in the initial inoculum and remained undetectable even after prolonged enrichment. *N. europaea* and *N. eutropha* were the dominant AOB across all enrichment stages, suggesting that prolonged cultivation was primarily shaped by the composition of the initial inoculum (Fig. 2a). From enrichment stage-I to III, *N. europaea* exhibited a slight increase from 5.1 × 10^5^ to 9.14 × 10^5^ copies/mg-biomass, whereas *N. eutropha* increased sharply from 8.43 × 10^3^ to 5.41 × 10^5^ copies/mg-biomass. *N. europaea* was already dominant in the initial inoculum, but continuous enrichment considerably increased *N. eutropha*. The *amoA* abundance of *N. eutropha* correlated positively with the nitrification rate (Pearson r = 0.72, p = 0.0001) shown in Fig. 1, indicating a greater contribution to ammonia oxidation than *N. europaea* (r = 0.19, p = 0.39). Given the strong adaptability to high ammonia environment, a consistent rise in *N. eutropha* abundance with 200 mg-N/L of 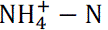 loading was not surprising (Fumasoli et al., 2015). Moreover, three-way ANOVA revealed that *amoA* gene abundance was strongly influenced by both enrichment age (partial η² = 0.63, p < 0.0001) and AOB species (partial η² = 0.57, p < 0.0001).

**Fig. 2.**
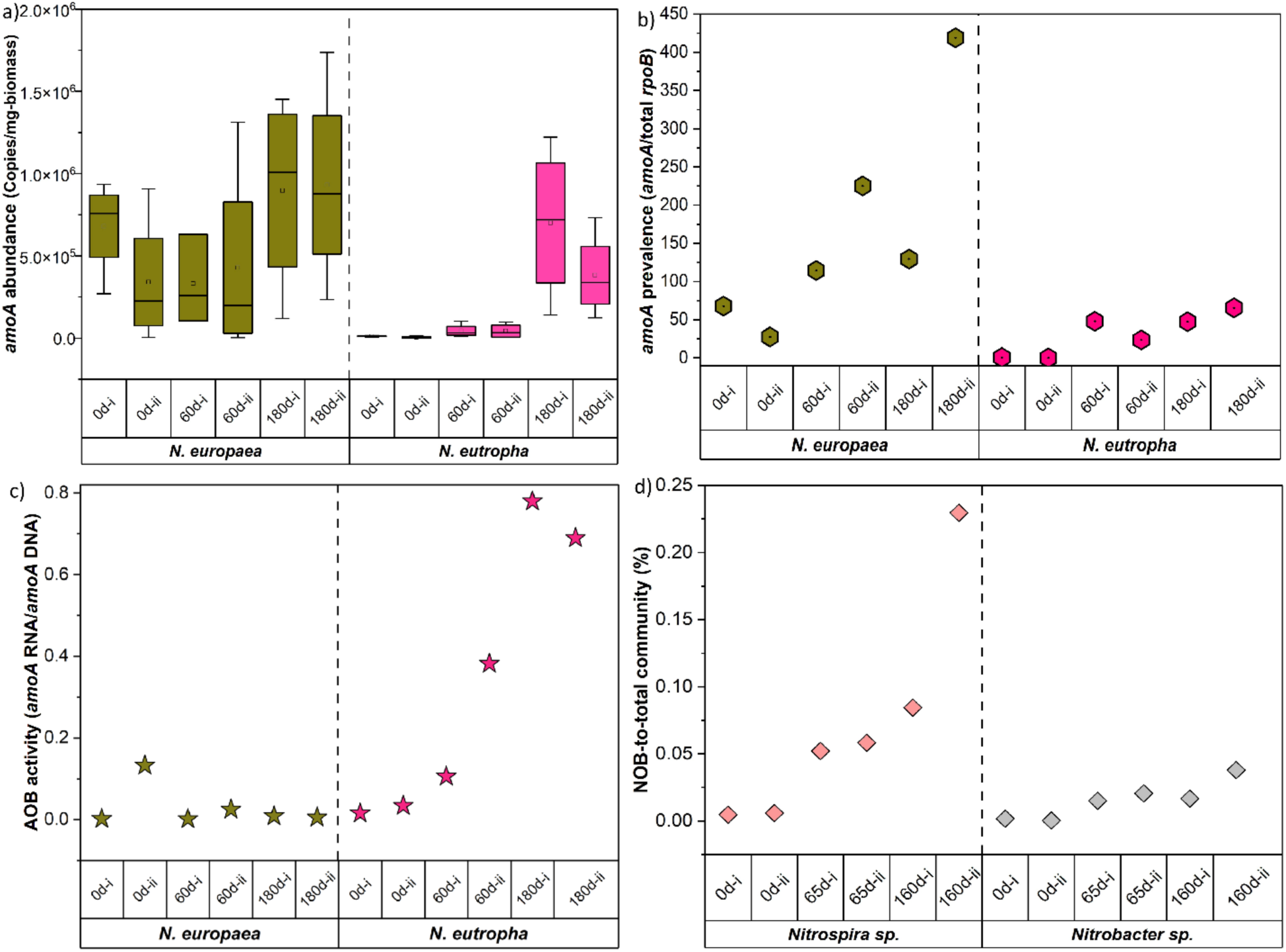
Profiles of AOB and NOB species across enrichment stages with biological replicates: (a) *amoA* gene abundance, (b) *amoA* gene abundance normalized to *rpoB*, (c) *amoA* transcriptional activity, (d) relative abundance of dominant NOB normalized to total 16S rRNA.

The community-normalized *amoA* abundance (*amoA*/*rpoB*) increased markedly over time for both AOBs (Fig. 2b). Although *rpoB* copy numbers declined from an average of ∼2.3 × 10^4^ copies/mg-biomass to 6.5 × 10^3^ copies/mg-biomass (from stage I to III), implying partial biomass washout. Whereas the higher *amoA*/*rpoB* indicate a functionally specialized community towards ammonia oxidation, driven by the prolonged ammonia loading. The synergistic coexistence of *N. europaea* and *N. eutropha* implies rapid and sustained ammonia-oxidizing capacity, consistent with the rising nitrification rates depicted in Fig. 1.

The ratio of relative abundance of *amoA* transcripts to *amoA* gene copies, representing transcriptional activity, was determined for both AOB species (Fig. 2c). Exposure to 200 mg-N/L of 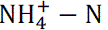 led to a progressive rise in *amoA* transcript levels with enrichment age. At 180d, the *N. eutropha* reached the highest transcription (∼7.3 × 10^−1^), whereas *N. europaea* remained substantially lower (∼6.6 × 10^−3^). It was interesting to note that despite the lower abundance of *N. eutropha* than *N. europaea*, the *amoA* genes were transcribed more intensely relative to its gene copy number across all enrichments. Although *N. europaea* and *N. eutropha* showed relatively high PCC with the AOR (r = 0.82 and 0.72, respectively), neither relationship was found statistically significant. Nevertheless, *N. eutropha* likely retained the dominant role in overall ammonia oxidation demonstrated by increased *amoA* transcription. Thereafter, a well acclimatized AOB consortium (∼180 d of cultivation period) was considered suitable for QS and QQ testing.

#### 3.1.2. Nitrite oxidation increased 8-fold during the enrichment period, resulting in a mixed *Nitrospira* and *Nitrobacter* NOB community

As shown in Fig. 1 and Fig. S1b, a minimal decrease in nitrite concentration without a corresponding rise in nitrate level during stage I indicated that the fresh inoculum contained a low fraction of active NOB, a condition that persisted for roughly 30 days. The concentrations of free nitrous acid remained ≤ 0.004 mg HNO₂-N/L, which was well below reported inhibitory thresholds for NOB (i.e. ∼0.02 mg HNO₂-N/L) (Owaes et al., 2023). With continued dosage of 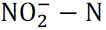, NOB growth and activity were progressively enhanced, leading to a marked increase in nitrite oxidation rate (NOR) from 0.6 mg-N g^−1^VSS h^−1^ (stage I) to 3.8 mg-N g^−1^VSS h^−1^ (stage II), which further increased to 4.8 mg-N g^−1^VSS h^−1^ by stage III (*p*<0.05). By day 160, the NOB bioreactor achieved stable nitrite oxidation (> 90% within 48 h) with a nitrite loading of 200 mg-N/L. As observed for AOB, prolonged enrichment substantially enhanced nitrate production compared with fresh sludge, confirming that increased NOB abundance and activity collectively determine the functional contributions within the nitrogen cycle (Zhang et al., 2023).

Quantitative analyses identified *Nitrospira* and *Nitrobacter* as the dominant NOB genera, consistent with earlier reports from wastewater-treatment systems (Shourjeh et al., 2021). Initially, *Nitrospira* and *Nitrobacter* were present at approximately 1.2 × 10^3^ and 2.3 × 10^2^ copies/mg-biomass, respectively, corresponding to 0.004–0.006 % and 0.0004–0.002 % of the total bacterial community, as quantified by NOB 16S rRNA gene abundance relative to total 16S rRNA gene quantified using universal primers. After 160 days of enrichment, their abundances increased to about 5.9 × 10^3^ and 1.1 × 10^3^ copies/mg-biomass (Fig. 2d), representing roughly four-fold enrichment. Although 16S rRNA gene copies of the overall community decreased by nearly an order of magnitude during enrichment (2.3 × 10^7^ and 4.6 × 10^6^ copies/mg-biomass), absolute NOB abundance continued to increase, confirming active proliferation and selective enrichment rather than a relative increase caused by biomass loss.

*Nitrospira* remained the predominant NOB genus, accounting for 0.08–0.23 % of the total community, although the overall NOB fraction was still lower than the levels generally reported in fully acclimated nitrifying systems (1 – 5% of the total bacterial community) (Yao and Peng, 2017). Both *Nitrospira* and *Nitrobacter* showed a strong positive correlation with NOR (r = 0.80 and 0.91, respectively), confirming their functional roles in nitrite oxidation. The observed predominance of *Nitrospira* in this study likely reflects its higher substrate affinity and adaptation to nitrite concentrations in the system (Fig. S2) (Wang et al., 2016). At stage III, the NOB activity was considered stable for subsequent tests.

### 3.2. AHL mixtures were determined based on abundance in the enriched sludge and on structural properties

Various nitrifying genera have been recognized as AHLs producers, and supplementation of exogenous AHLs has been shown to enhance nitrification efficiency in several biological systems (Sun et al., 2018; Yang et al., 2024; Zhao et al., 2025). Given the prevalence and diversity of these autoinducers, the AHL profile of the enriched sludge was first analyzed and categorized according to abundance. AHLs identified at high-abundance (C6-, C8-, C10-HSL), medium-abundance (3OC6-, 3OC10, 3OC12-HSL), and low-abundance (C4-, 3OC8, C12-HSL) were quantified at 22.7±4.2, 13.7±4.5, and 3.2± 0.3 µmol/L, respectively. Selective enrichment of nitrifying species reduced overall biomass (Figure S3), potentially lowering the proportion of heterotrophic AHL-producing bacteria. Moreover, not all nitrifiers are known for AHLs production (Wang et al., 2021). To examine how different naturally occurring AHL groups influence nitrification, enriched AOB-and NOB-sludge were spiked with AHL mixtures (to 500 nmol/L) representing different abundance categories low-(C4-, 3OC8-, C12-HSL), medium-(3OC6-, 3OC10-, 3OC12-HSL), high-abundance (C6-, C8-, C10-HSL), all identified AHLs, and based on structural classes (short-chain, long-chain, oxo-acyl, and acyl) (Table S2). As QS and QQ coexist and jointly regulate nitrogen removal (Feng et al., 2019), a well-reported AHL-quenching enzyme acylase I was also applied to examine the effect of QQ on nitrifying activity of AOB and NOB enrichments (Yu et al., 2018).

#### 3.2.1. AHLs enhanced ammonia oxidation, demonstrating modest trends related to type of molecule

The enrichment experiment demonstrated increased nitrifying activity in AOB sludge associated with cell density. The potential role of AHLs in regulating microbial density was assessed in detail as illustrated in Fig. 3a. As expected, relative to the fresh control (CnF, ≈ 2.9 mg-N g⁻¹ VSS h⁻¹), the enriched control (CnEn) achieved about 1.8-fold higher ammonia oxidation rates (≈ 5.3 mg-N g⁻¹ VSS h⁻¹), consistent with greater *amoA* abundance and transcription capacity in both *N. eutropha* and *N. europaea* (Fig. 4-a2, S4).

**Fig. 3.**
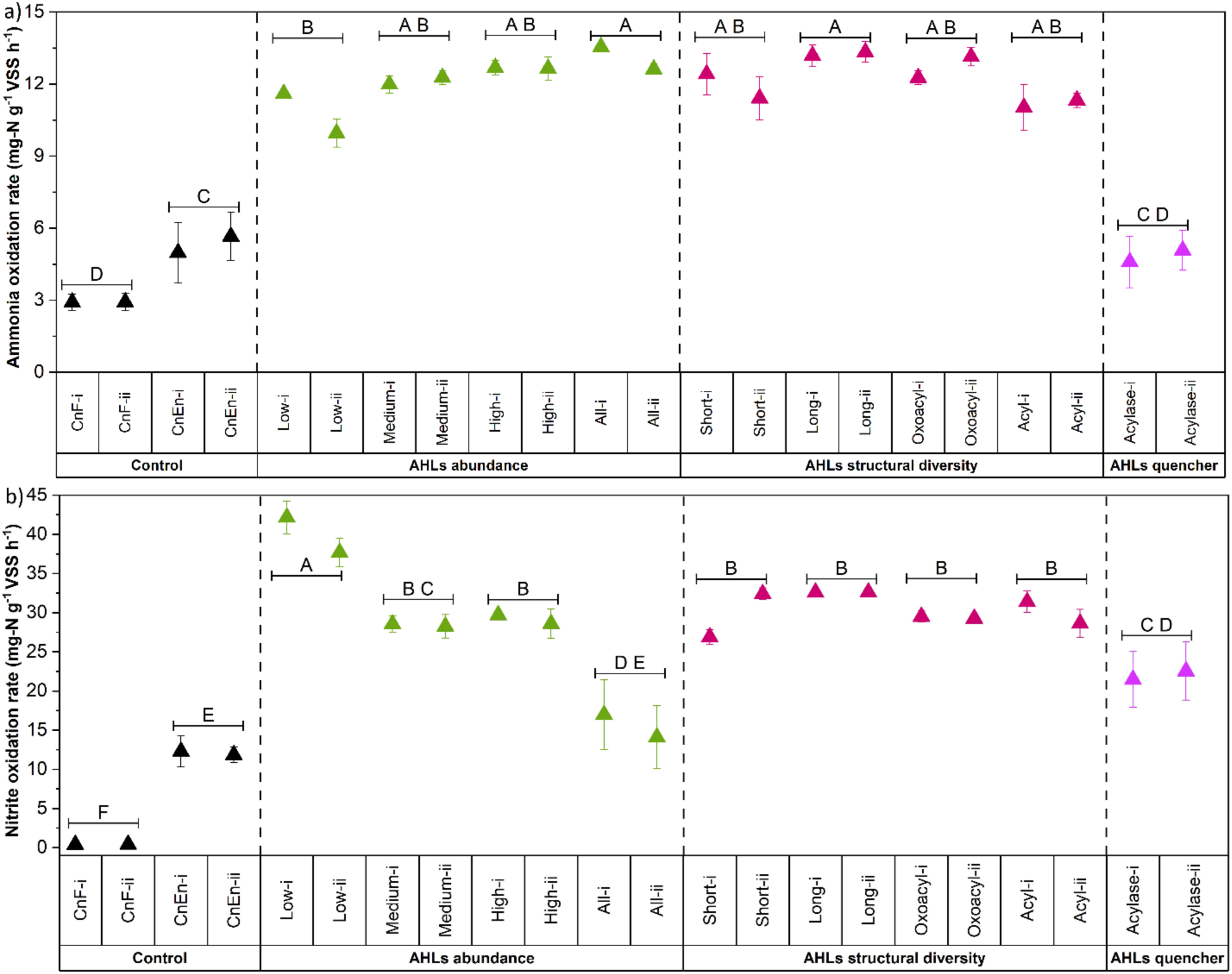
AHLs influence on: (a) Ammonia oxidation rate in AOB sludge, (b) Nitrite oxidation rate in NOB sludge measured by a linear fit. Groups not sharing the same letter (A-F) differ significantly as per Tukey’s test (*p*<0.05).

**Fig. 4.**
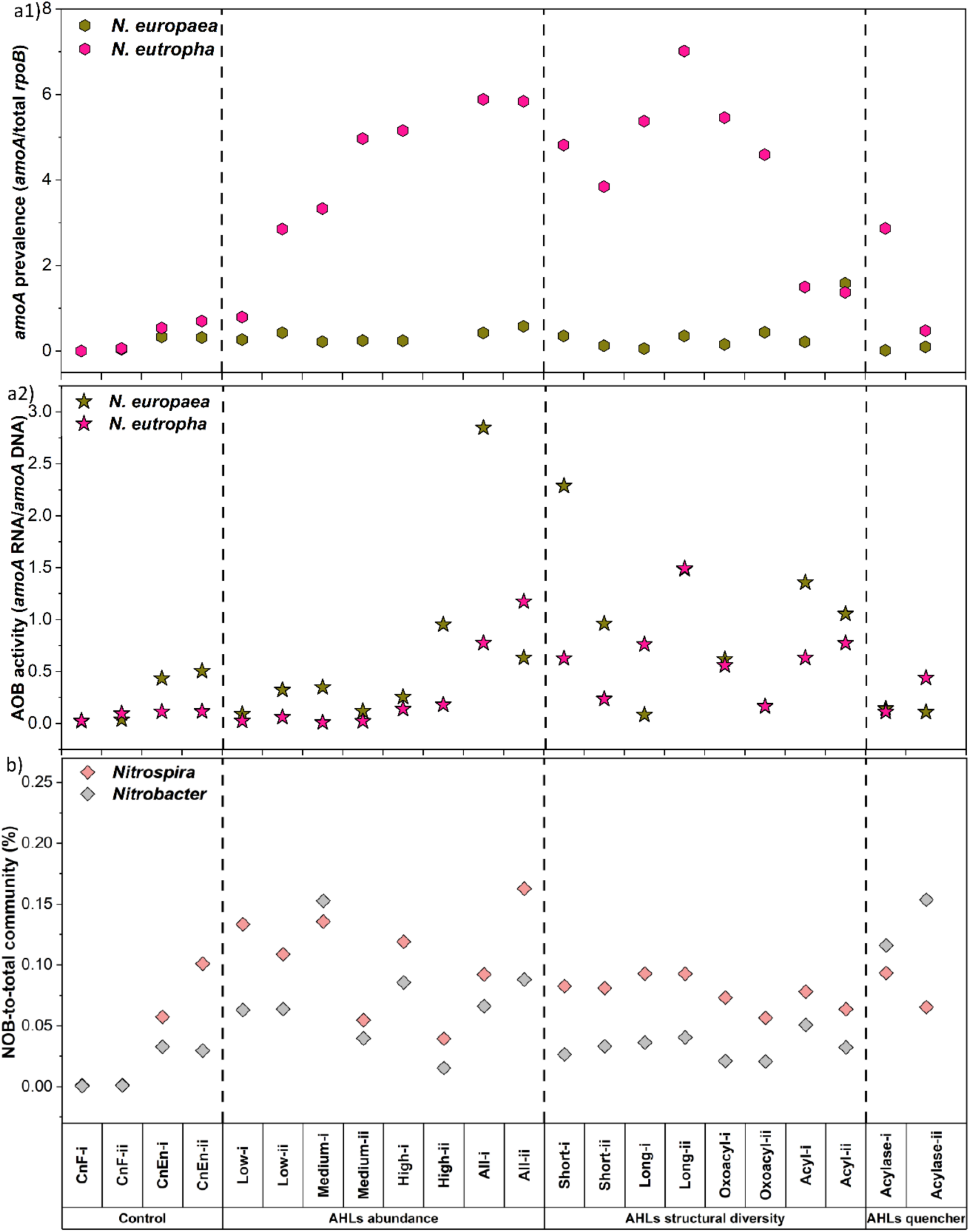
AHLs influence across treatments with biological replicates: (a1) *amoA* gene abundance normalized to *rpoB* at 4-hour timepoint, (a2) *amoA* transcriptional activity at 12-hour timepoint, (b) relative abundance of dominant NOB normalized to total 16S rRNA.

Exogenous addition of AHLs further enhanced AOR by 2 to 2.5-fold over the enriched control (CnEn), and 3.7 to 4.5-fold over the fresh sludge (CnF). The AOR demonstrated moderate increases with AHL abundance, from 10.8±1.2 mg-N g^−1^VSS h^−1^ for the low-abundance set to 13.1±0.7 mg-N g^−1^VSS h^−1^ when all AHLs were supplemented. This trend reiterates that AHLs that were present at higher concentrations positively influenced ammonia oxidation activity. Of note, the High-abundance and All-AHLs treatments yielded similar rates (12.7 vs 13.1 mg-N g^−1^VSS h^−1^, *p*>0.05), even though the latter contained three-times more total signal load (4500 nmol/L) compared to former (1500 nmol/L), which could indicate a near-saturation response to quorum cues. Future work could elucidate how AHL saturation occurs in mixtures relative to individual compounds. Given the signaling threshold and functional plateau, we inferred that the dominant AHL set (C6-, C8-, C10-HSL) appeared to be the most efficient signals for stimulating oxidation. Among structural variants, long-chain and Oxoacyl sets supported the highest AOR (13.3 and 12.7 mg-N g^−1^VSS h^−1^, respectively), though again the effect was modest compared to other structural variants. Long chain AHLs generally exhibit higher hydrophobicity; their increased hydrophobicity and persistence likely prolonged signal retention within sludge flocs, strengthening and sustaining quorum activation (Chen et al., 2019; Liu et al., 2022). This shows that AHLs with extended acyl chains and substituent chemistry play a crucial role in signal persistence and bioavailability, ultimately influencing nitrifying activity.

In contrast, AHL quenching via Acylase lowered AOR to 4.8±0.3 mg-N g^−1^VSS h^−1^, below the rate of CnEn but above the CnF. Quorum quenching coincided with smaller floc particle size (D₉₀ ≈ 123 µm vs ≈ 357 µm in AHL-treated sludge), which could improve substrate diffusion and partly offset the loss of QS-mediated stimulation. The smaller flocs are considered favorable for nitrifiers due to lower diffusional resistance for both nitrogen substrate and oxygen (Britschgi et al., 2025). Collectively, these trends confirm that AHL signaling augments oxidation efficiency, while biomass enrichment and substrate accessibility also modulate overall nitrification performance.

Parallel molecular analyses were conducted to evaluate correlation with rates and functional changes. AOB species increased in *amoA* gene abundance relative to fresh control, but *N. eutropha* remained numerically dominant throughout all treatments (Fig. S4), aligning with the corresponding rise in AOR. Across AHL spiking sets, the relative abundance of *N. eutropha* as measured by the *amoA*/*rpoB* ratio progressively increased from the low– to the All-AHL group and correlated with the AOR (r = 0.80, *p*<0.01), whereas *N. europaea* showed a weaker and non-significant relationship (r = 0.32, *p*>0.05) (Fig. 4-a1). Within the structural sets, long-chain and oxoacyl AHLs exhibited modestly higher *amoA/rpoB* values, reinforcing their potential impact on nitrification.

As expected, during the 36-h oxidation time, the abundances of *amoA* DNA remained relatively stable across AHL treatments. This suggests that, over the time scales of these experiments, AHL supplementation primarily influenced ammonia oxidation activity and observed rate patterns stemmed from transcriptional regulation rather than *amoA* community abundance. Previous studies also reported no noticeable impact of AHLs addition on nitrifying community composition under steady-state conditions (Zeng and Hu, 2023). The *amoA* abundance of both AOBs remained stable during the Acylase treatment, as the enzyme mainly interrupts the QS pathway that regulates transcription via AHL degradation, with minimal impact on population growth. The reduction in AOB activity rather than a decrease in abundance observed under Acylase treatment was consistent with the previous report (Yu et al., 2018).

Correspondingly, the expression of *amoA* (*amoA*-RNA/*amoA*-DNA ratios) increased under AHL exposure, with a positive correlation between the level of *amoA* transcripts and AOR at 12h for both AOBs (r = 0.45, *p*<0.05) (Fig. 4-a2). During the active oxidation phase (12h), accumulated AHLs likely reached levels sufficient to activate the receptor protein, which in turn triggered *amoA* gene expression. AHLs spiking based on structural diversity resulted in distinct levels for *amoA* expression in contrast to other groups. *N*. *eutropha* displayed particularly strong activation in the long-chain AHL group with *amoA* RNA of 5.01× 10^5^ copies/mg-biomass and RNA/DNA ratio of 1.13, confirming its higher transcriptional responsiveness to QS signals. Transcriptional activity under acylase treatment remained comparable to some AHL-spiked sets, suggesting incomplete quenching and demonstrating that the AOB sludge retained considerable ammonia-oxidizing potential and functional resilience even under partial QS suppression.

In summary, AHLs spiking significantly boosted ammonia oxidation and *amoA* transcription, predominantly through activation of *N. eutropha*. Higher AOR corresponded with elevated *amoA*/*rpoB* ratios and increased *amoA* transcripts under specific AHL groups, confirming the pivotal role of quorum signaling in regulating AOB metabolic activity.

#### 3.2.2. The type of AHL molecule had more pronounced impact on NOB sludge, compared with AOB sludge

One-way ANOVA indicated a significant difference in nitrite oxidation between CnF and CnEn groups, reiterating the influence of higher NOB biomass density (*p*<0.05; Fig. 3b). Exogenous AHL addition showed a distinct response in NOR across the molecule types; the low-abundance AHL set yielded the highest NOR of 40 ± 3.2 mg N g⁻¹ VSS h⁻¹, approximately 3.3 times greater than the enriched control (12 mg N g⁻¹ VSS h⁻¹). However, oxidation declined progressively under the medium-, high-, and all-AHL treatments, indicating differing sensitivities of the NOB guild to specific signal types and levels. It was interesting to note that the lowest proportion of AHLs in AOB sludge resulted in the fastest NOR in NOB enriched consortium. Because the AHL abundance categories were derived from the AOB enriched sludge, the inverse trend observed here highlights the contrasting impact of AHLs on the two nitrifying guilds: ones beneficial to AOB appeared inhibitory to NOB. The decline in nitrite oxidation at elevated total AHL concentrations (4,500 nmol/L in the All-AHL set), was likely owing to formation of larger, oxygen-limited flocs that restricted nutrient diffusion, resulting in reduced NOB activity (Britschgi et al., 2025). When AHLs were grouped by structural class, NOR did not differ markedly among short-, long-, oxoacyl-, or acyl-chain sets, though all treatments exceeded the controls (*p*<0.001).

Contradictory to AOB, QQ through acylase enhanced NOB activity rather than suppressing it. The NOR roughly doubled relative to the enriched control, coinciding with smaller and less cohesive flocs (∼ 100 µm) that possibly improved mass transfer of oxygen and nitrite. Moreover, disruption of QS may also have diminished intraguild competition from heterotrophic populations within the sludge, further facilitating NOB oxidation. These observations suggest that nitrite oxidation activity in NOB is primarily governed by substrate and oxygen accessibility rather than intercellular signaling cues.

Parallel NOB community analyses during the period of maximal nitrite oxidation (∼6h) revealed that *Nitrospira* remained the dominant genus across most treatments (Fig. 4b). Both *Nitrospira* and *Nitrobacter* increased in relative abundance from the fresh to the enriched controls, and their combined copy numbers correlated strongly with NOR in the control treatments (PCC = 0.98). Across the AHL additions, NOR variations were not directly linked to NOB abundance, implying that nitrite oxidation was modulated through signaling interference at the functional rather than population shifts. Increases in NOR driven by low-abundance AHLs (C4-, 3OC8-, C12-HSL) suggest that these signal types promote metabolic coordination within NOBs. Conversely, a progressive decline under medium, high, and all groups indicates that certain AHLs likely exerted an inhibitory or competitive effects on NOB activity without substantially changing cell number. Within the AHL structural-diversity, 3OC8– and C12-HSL in the long-chain and C4– and C12-HSL in the acyl group yielded the strongest oxidation responses, indicating enhanced utilization of these compounds as signaling cues.

Under acylase-mediated quenching, *Nitrobacter* showed a relative increase compared with the control and AHL-treated systems, suggesting a compensatory functional shift. As *Nitrobacter* carries LuxI/LuxR homologs (Mellbye et al., 2017), it may up-regulate signal synthesis under QS-deficient conditions to restore communication and sustain oxidation activity under stress. Overall, the enhanced NOR alongside stable or increased NOB abundance during quenching treatments indicates that the improved performance was not due to biomass loss but to changes in community interactions and diffusion conditions. Collectively, these results demonstrate that AHL supplementation and quenching differentially modulate NOB physiology: low-abundance AHLs can stimulate nitrite oxidation, whereas elevated or mixed AHL loads inhibit it, and QS disruption through acylase promotes functional resilience by improving mass transfer and shifting NOB dominance from *Nitrospira* to *Nitrobacter*.

### 3.3. AI-2 & selected quenchers for nitrifying sludge

The effects of AI-2–mediated QS and its quenching by quercetin on the nitrifying activity of AOB and NOB consortia were evaluated, along with the combined impact of AHL and AI-2 quenchers (quercetin + acylase) on overall nitrification rates.

#### 3.3.1. AI-2 supplementation reduced AOR, but quenching AI-2 had no effect

Fig. 5a demonstrated the comparative effect of spiking and quenching AI-2 on AOB enriched sludge. AOR declined from 5.3 mg-N g⁻¹ VSS h⁻¹ in the enriched control (CnEn) to 2 mg N g⁻¹ VSS h⁻¹ after AI-2 addition, implying that excessive AI-2 perturbed the nitrifier metabolism and suppressed AOB activity. This reduction coincided with a near two-fold increase in average floc size, consistent with reports that AI-2/LuxS signaling promotes biofilm and granule formation (Hu et al., 2022). Larger, denser flocs likely restrict oxygen diffusion, thereby diminishing the accessible surface area for ammonia oxidation.

**Fig. 5.**
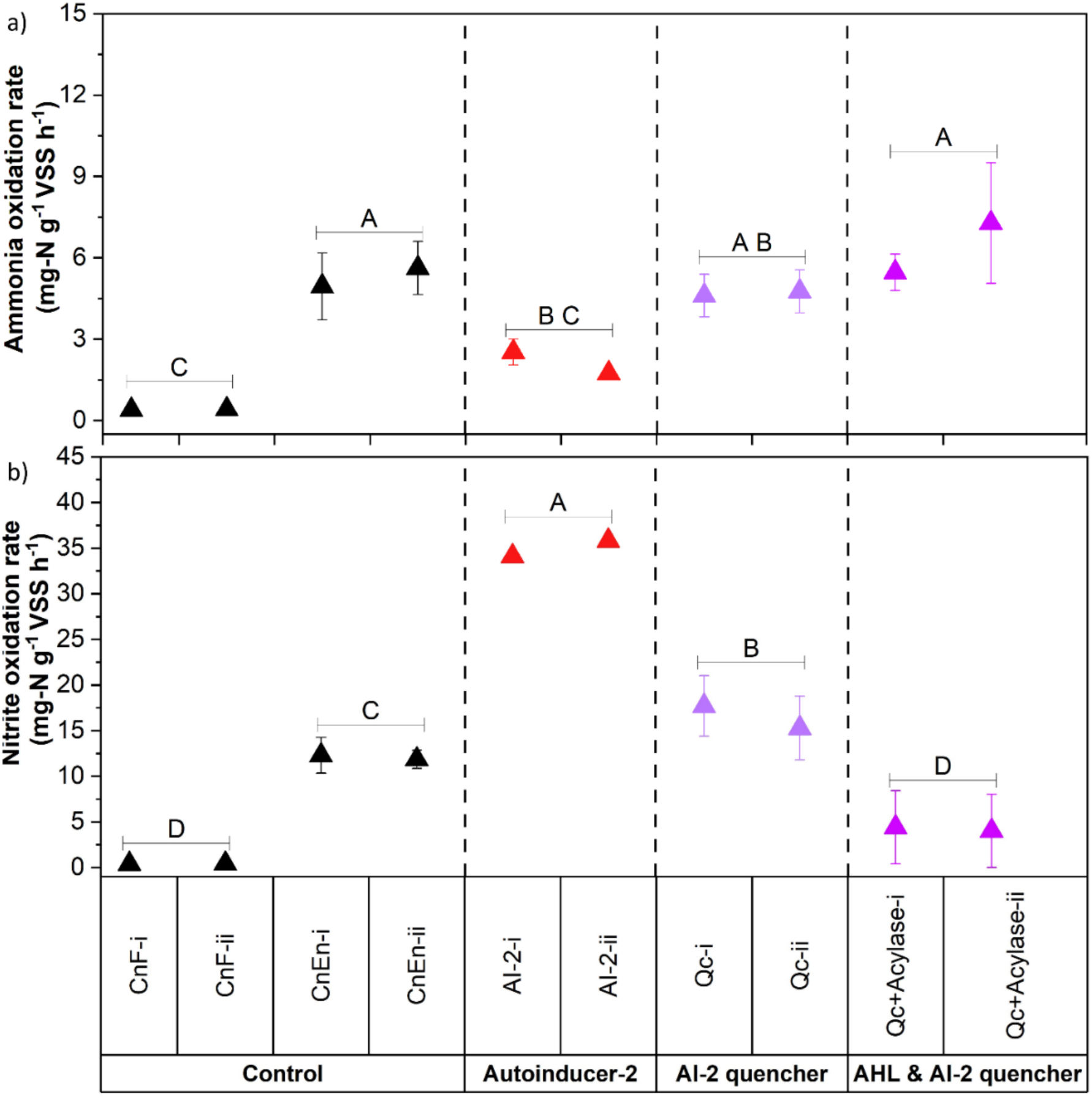
AI-2 influence on: (a) Ammonia oxidation rate in AOB sludge, (b) Nitrite oxidation rate in NOB sludge measured by a linear fit. Groups not sharing the same letter (A-F) differ significantly as per Tukey’s test (*p*<0.05).

In contrast, a notably higher AOR was observed under quenching conditions (Qc, Qc+Acylase), suggesting that suppression of signaling favored ammonia oxidation. Both quercetin and combined quenching with quercetin + acylase increased AOR reaching 4.7 ± 0.1 mg-N g⁻¹ VSS h⁻¹ and 6.4 ± 1.3 mg −N g⁻¹ VSS h⁻¹, respectively. This reflects that suppression of AI-2 signaling was primarily contributing to the 70 % increase in activity, while simultaneous removal of both AI-2 and AHLs produced a modest additional gain. Smaller, less cohesive flocs observed in the quenched reactors shortened diffusion paths enabling more efficient oxygen and substrate penetration, further augmenting AOR.

The variation in AOR among treatments closely followed changes in abundance for the two dominant AOBs (Fig. S5). AI-2 exposure reduced *amoA* gene copies in both organisms, but the decline was more pronounced and persistent for *N. eutropha*. Quenching restored *amoA* abundance, with a more obvious rebound of *N. eutropha*, consistent with the elevated oxidation rates. *N. europaea* appeared to be less dynamic under sensing and quenching of AI-2, implying lower dependence on AI-2 cues. The coexistence of both populations with a contrasting response demonstrates functional redundancy within the AOB consortium. Wherein, *N. europaea* maintains baseline oxidation while *N. eutropha* provides responsiveness and resilience to changes in signaling intensity, allowing a community to sustain performance under diverse signaling disturbances.

The housekeeping gene *rpoB* remained relatively stable (2 – 4 × 10⁴ copies mg⁻¹ biomass) across all treatments, differing by less than two orders of magnitude (Fig. 6-a1). This confirms that total bacterial biomass in the nitrifying sludge was largely stable and that no significant biomass wash-out occurred during AI-2 stimulation or quenching. Hence, the differing responses observed under AI-2 perturbation for both AOBs were driven by functional regulation.

**Fig. 6.**
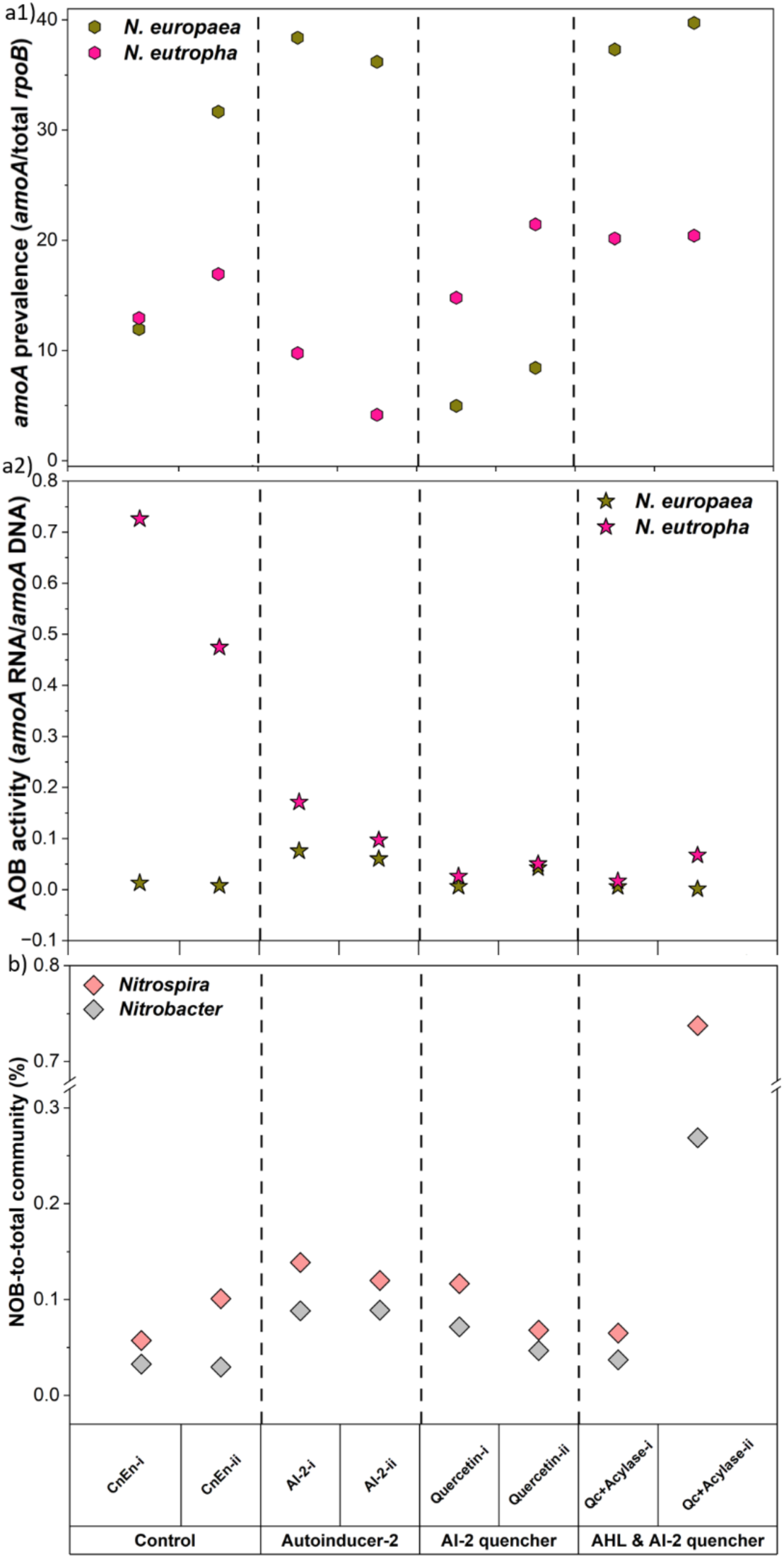
AI-2 and quenching role across treatments with biological replicates: (a1) *amoA* gene abundance normalized to *rpoB* at 4-hour timepoint, (a2) *amoA* transcriptional activity at 12-hour timepoint, (b) relative abundance of dominant NOB normalized to total 16S rRNA.

Notably, AHL and AI-2 exerted contrasting effects on the two AOB populations. Whereas AHL supplementation stimulated *N. eutropha*, increasing the *amoA*/*rpoB* ratio, AI-2 exposure decreased its relative abundance (Fig. 6-a1). In contrast, *N. europaea* was largely unresponsive to AHL supplementation yet displayed increased *amoA* abundance when exposed to elevated AI-2. This divergence highlights that the two signaling pathways distinctly shape AOB physiology and contribute differently to overall nitrification performance.

Consistent with this, transcriptional analyses clarified the regulatory effects of AI-2 (Fig. 6-a2). *N. europaea* exhibited nearly a ten-fold rise in *amoA* transcript ratio (≈ 0.07 vs 0.01 in control) when exposed to excess AI-2, whereas *N. eutropha* transcription decreased sharply from 0.6 to 0.1. During AI-2 quenching, transcript levels of *N. europaea* fell to near-control values, and *N*. *eutropha* partially recovered (≈ 0.03–0.05), while the combined Qc + Acylase treatment equalized both species at low, stable transcriptional states. Overall, AI-2 manipulation altered nitrification primarily through regulatory rather than biomass effects. Over-stimulation by AI-2 created metabolic imbalance between the two AOB species, whereas controlled quorum-signal quenching restored balanced *amoA* expression and higher cumulative oxidation. These findings underscore the dual nature of AI-2 in nitrifying consortia as both a coordination signal and a potential inhibitor when present above its physiological threshold.

#### 3.3.2. AI-2 supplementation increased NOR substantially

In contrast to the inhibitory effect on AOB, AI-2 spiking significantly enhanced NOR compared to control and quenching conditions (*p*<0.001) (Fig. 5b). AI-2 addition increased the NOR from 12 mg-N g⁻¹ VSS h⁻¹ in the enriched control (CnEn) to 36 mg-N g⁻¹ VSS h⁻¹ (*p* < 0.001), representing nearly a threefold enhancement. This strong stimulation suggests that AI-2–driven interspecies communication promoted collaborative metabolism among NOBs, likely by improving cell–cell coordination and expression of nitrite-oxidizing genes. During quercetin-based AI-2 quenching, the NOR was 16 mg-N g⁻¹ VSS h⁻¹, comparable to the CnEn. This suggests that AI-2 signaling transiently stimulated NOB activity, but its suppression simply reverted the system to its baseline without further inhibition. However, the synergistic action of quercetin and acylase (Qc + Acylase) drastically curtailed NOB activity, reducing NOR by 89 % relative to the AI-2-supplemented set. This outcome demonstrates that simultaneous quenching of AI-2 and AHL cues exerts a strong inhibitory effect on nitrite oxidation. Given the contrasting signaling requirements, physiological, and metabolic properties of AOB and NOB guilds, such opposite responses to AI-2 and AHL manipulation on nitrification performance were expected and are consistent with their distinct ecophysiology.

Correlation analysis (Pearson’s r) further supported these observations. NOR correlated more strongly with *Nitrospira* (r = 0.76) than with *Nitrobacter* (r = 0.56), emphasizing the dominant contribution of *Nitrospira* to nitrite oxidation under all conditions (Fig. 6b). AI-2 supplementation increased the relative abundance of both genera, with *Nitrospira* rising from 0.08 % to 0.13 % and *Nitrobacter* from 0.03 % to 0.09 %. These increases confirm that AI-2 signaling enhanced NOB proliferation and functionality.

When AI-2 was quenched, NOR and *Nitrobacter* fractions decreased toward the control level, whereas *Nitrospira* remained comparatively stable, potentially indicating that disruption of AI-2 primarily affected the *Nitrobacter*-associated pathways supporting rapid oxidation. This compositional shift likely contributed to reduced nitrite oxidation, as *Nitrospira* remained relatively abundant. Under dual quenching (Qc + Acylase), NOR dropped further despite detectable NOB abundances, showing that the inhibitors acted mainly through metabolic rather than compositional suppression.

Taken together, these results demonstrate that AI-2 serves as a positive regulatory cue for NOBs, in sharp contrast to its inhibitory role for AOBs. AI-2 enrichment fosters cooperative behavior and higher oxidation rates through strengthened communication among NOB species, whereas quenching, especially coupled quenching of AHL and AI-2, interrupts these interactions and reduces nitrite-oxidizing efficiency. The findings highlight the opposing roles of QS signals within the two nitrifying guilds and suggest that optimal nitrification depends on a balanced level of interspecies communication.

## 4. Conclusions

The study highlights distinct regulatory roles of AI-2/AHL-mediated communication within the nitrifying consortium and its potential to decouple the two oxidation steps when artificially perturbed. Manipulation of AHLs revealed guild-specific signaling effects. AHL supplementation enhanced ammonia oxidation, primarily through transcriptional activation of *amoA* in *N. eutropha*, whereas excessive or mixed AHLs inhibited nitrite oxidation. Correspondingly, AHLs quenching with acylase reduced but did not eliminate AOB activity and improved NOB performance by alleviating diffusion limitations and shifting community dominance toward *Nitrobacter*.

Elevated AI-2 levels disrupted AOB metabolism, enlarged sludge flocs, and suppressed ammonia oxidation, whereas signal quenching, particularly the combined removal of AI-2 and AHLs restored balanced *amoA* expression and resulted in the highest overall AOR. In contrast, AI-2 stimulated NOB activity and proliferation, confirming its role as a positive cue for nitrite oxidation. Dual quenching of AI-2 and AHL signals curtailed NOB performance, highlighting the contrasting regulatory requirements of the two nitrifying guilds. This finding has wider implications for effective employment in settings where suppressed NOB has an overall advantage in nitrification. Taken together, the results demonstrate that quorum signals govern nitrification primarily through functional regulation. AHLs and AI-2 act as complementary yet opposing controls for the AOB and NOB populations and maintaining these signaling molecules within an optimal physiological window is crucial for synchronized oxidation of ammonia and nitrite and for maximizing overall nitrogen removal efficiency in biological treatment systems.

## Acknowledgements

This material is based upon work supported by the National Science Foundation under a CAREER Award (2349328) and an Understanding the Rules of Life award (Award 2319124). This work was also supported in part by the Engineering Research Centers Program of the National Science Foundation under NSF Cooperative Agreement No. EEC-2133504.

## Declaration of AI-assisted technologies in the manuscript preparation process

During the preparation of this work the authors used ChatGPT and DukeGPT to improve language clarity, and readability. All content assisted by these tools was carefully reviewed and edited by the authors who take full responsibility for the content of the published article.

