## Supplemental Information for "Autoinducer-2 and acyl homoserine lactones have contrasting effects on ammonia and nitrite-oxidizing sludge"

### Supporting Information

#### 1.1. Composition of synthetic feed

For ammonia oxidizing bacteria (AOB) and nitrite oxidizing bacteria (NOB) enrichments, $(NH_4)_2SO_4$ and $NaNO_2$ with a concentration of 944 mg/L and 986 mg/L (with N loading of 200 mg/L) were used as sole nitrogen source, respectively. The remaining composition of the synthetic feed was identical for both enrichments as described in Table S1.

Table S1: Synthetic feed composition for bioreactors.

| **Feed solution** | **mg/L** | **Trace element solution** | **mg/L** |
|---|---|---|---|
| $NaHCO_3$ | 1150 | $Na_2EDTA$ | 0.88 |
| $MgSO_4.7H_2O$ | 6.6 | $Na_2MoO_4.2H_2O$ | 0.01 |
| $MgCl_2.6H_2O$ | 5 | $MnSO_4.H_2O$ | 0.045 |
| $CaCl_2.2H_2O$ | 6 | $CoCl_2.7H_2O$ | 0.0004 |
| $K_2HPO_4$ | 16 | $ZnCl_2$ | 0.06 |
| $KH_2PO_4$ | 5 | $CuSO_4.5H_2O$ | 0.255 |
| | | $FeSO_4.7H_2O$ | 1.1 |
| | | $H_3BO_3$ | 0.13 |

Table S2: AHLs categorized for the batch tests.

| Set | Treatment |
|---|---|
| 1. Control with fresh sludge (CnF) | No autoinducers added |
| 2. Control with enriched sludge (CnEn) | No autoinducers added |
| 3. Low abundant AHLs (Low) | C4-, 3OC8-, C12-HSL |
| 4. Average abundant AHLs (Medium) | 3OC6-, 3OC10-, 3OC12-HSL |
| 5. Dominant AHLs (High) | C6-, C8-, C10-HSL |
| 6. Mixture of 9 AHLs (All) | C4-, C6-, 3OC6-, C8-, 3OC8-, C10-, 3OC10-, C12-, 3OC12-HSL |
| 7. Short-chain AHLs (Short) | C4-, C6-, 3OC6-, C8-HSL |
| 8. Long-chain AHLs (Long) | 3OC8-, C10-, 3OC10-, C12-, 3OC12-HSL |
| 9. AHLs with oxo functional group (Oxoacyl) | 3OC6-, 3OC8-, 3OC10-, 3OC12-HSL |
| 10. AHLs with no functional group (Acyl) | C4-, C6-, C8-, C10-, C12-HSL |
| 11. AHLs quencher (Acylase) | Acylase I (EC 3.5.1.14) |

#### 1.2. AHLs extraction and detection through UHPLC-HRMS

Sludge supernatant and ethyl acetate were mixed (1:1) vigorously, and the organic layer was collected using a separatory funnel, dehydrated using anhydrous sodium sulfate, and dried via SpeedVac (Thermo Savant, SPD121P). The residue was re-dissolved in 1 mL of ethyl acetate and transferred to glass vials. The sample was slowly blow-dried via nitrogen ($N_2$); the remaining residue was dissolved in 100 µL of acetonitrile; filtered using 0.22 µm PTFE membranes and stored at -20°C. Samples were concentrated 1000 times and quantified by mass spectrometry with a 7-point internal calibration curve ranging from 0 to 1000 nmol/L; using C7-HSL, C4-HSL-d5, and C6-HSL-d3 as internal standards.

Sample analysis was performed by ultra-high pressure liquid chromatography-high resolution mass spectrometry (UHPLC-HRMS) using an Orbitrap Fusion Lumos MS and Vanquish Flex MS (both from ThermoFisher Scientific). During analysis, samples were held at 4°C in the autosampler and a 10 µL injection was separated using a C18 column (Hypersil Gold, with length and diameter of 100 and 2.1mm respectively, 1.9µm ThermoFisher Scientific, USA) with 0.1% formic acid (A) and acetonitrile with 0.1% formic acid (B) as the mobile phase at a gradient of 5-100% within 15 min at a flow rate of 0.25 mL/min and a column temperature of 30°C using a Vanquish UHPLC system. Analytes were detecting using an Orbitrap Fusion Lumos Tribrid HRMS in positive ion mode and the instrument settings were set as follows: cation spray voltage, 3.5kV; sheath gas, 30 (arbitrary units); auxiliary gas, 10 (arbitrary units); ion transfer tube and sprayer temperature, 300 °C. Analytes were detected using a TSIM method using the ion trap detector and processed using Thermo TraceFinder 5.1 (ThermoFisher Scientific).

**1.3. AI-2 detection through UHPLC-HRMS**

Sample analytes were quantified using the same LC-HRMS as described for AHLs. UHPLC separation occurred after a 10 µL injection onto the C18 column and separated with a flow rate of 0.3 mL/min and run time of 20 min, with at 5 min equilibration before the run. Mobile phases of water (A) and acetonitrile (B) were used for separation and the gradient started at 5%B and gradually increased to 100%B over 18 minutes, then held at 100%B for 2 minutes. The instrument settings using Orbitrap detector were set as follows: cation spray voltage, 3.5kV; anionic spray voltage, 2.5kV; sheath gas, 30 Arb; auxiliary gas, 10 Arb; ion transfer tube and sprayer temperature, 325 and 300 °C, respectively.

**1.4. ddPCR reaction mixture and thermal cycling condition**

In 96-well plates, each well contains 22 µL reaction mixture of 5.5 µL of ddPCR Supermix for probes (Bio-Rad), 0.147 µL from mixture of forward/reverse primer (900 nM) and probe (250 nM), 0.27 µL of DTT (300 mM), and 14.08 µL of nuclease-free water followed by 2 µL of template. Of note, each plate had no template control (NTC) and a positive control (GeneBlock (IDT)from each amplicon). The plates were sealed, briefly vortexed, and centrifuged at 2500 rpm for 2 min. Droplets were subsequently generated using an automated droplet generator (Bio-Rad). The thermal cycler was performed with the manufacturers conditions including 95 °C for 10 min, 40 cycles of 94 °C for 30 sec, 2 min hold at the annealing temperature (Table S4), and 98 °C for 10 min. Annealing temperature for all primer/probe set was optimized by preliminary testing through temperature gradient between 54 ºC and 65 ºC, aiming for optimal separation between positive and negative droplets.

Table S3: List of primers/probe used in the study.

| Target organisms | Target genes | Primers/Probe | Annealing temp (ºC) | Reference |
|---|---|---|---|---|
| *N. eutropha* | amoA | F (TCC ACT CAA TTT TGT AAC CCC)<br>R (ATC AGG CCA AAG AAT CCA CC)<br>Probe (56-FAM/CAA CCA GTT /ZEN/ ACG TGT CAG ATA CAT TGT GAA ATC C/3LABkFQ) | 58.1 | (Orschler et al., 2020) |
| *N. europaea* | amoA | F (CAC ACT ACC CCA TCA ACT TC)<br>R (GTC CCA TGT AAT CAG CCA TC)<br>Probe used for DNA analysis (5HEX/ATA GAA CAG CAG ACC GAA GAA TCC ACC TCC AAC CA/3 BHQ) or<br>Probe used for RNA analysis (5HEX/TAG AAC AGC/ZEN/AGA CCG AAG AAT CCA /3IABkFQ) | 58.1 | (Orschler et al., 2020) |
| *Nitrosospira* | amoA | F (GGG GTT TCT ACT GGT GGT)<br>R (CCC CTC KGS AAA GCC TTC TTC)<br>Probe (5Cy5/CCG ACS CAC/TAO/CTG CCG CTG G/3IAbRQSp) | 58.1 | (Kim et al., 2008) |
| Total | rpoB | F (CGA ACA TCG GTC TGA TCA ACT C)<br>R (GTT GCA TGT TCG CAC CCA T) | 56 | (Silkie & Nelson, 2009) |
| *Nitrobacter* | 16S rRNA | F (ACC CCT AGC AAA TCT CAA AAA ACC G)<br>R (CTT CAC CCC AGT CGC TGA CC)<br>Probe (5SUN/GGT CGG CTG /ZEN/CCT CCC TTG CGG GTT /3IABkFQ) | 62.7 | Modified from: (Graham et al., 2007) |
| *Nitrospira* | 16S rRNA | F (GCG GTG AAA TGC GTA GAK ATC G)<br>R (TCA GCG TCA GRW AYG TTC CAG AG)<br>Probe (56-FAM/CGC CGC CTT /ZEN/ CGC CAC CG/3IABkFQ) | 62.7 | Modified from: (Graham et al., 2007) |
| Total | 16S rRNA | 16S_1055 F (ATG GCT GTC GTC AGC T)<br>16S_1392 r (ACG GGC GGT GTG TAC) | 55 | (Harms et al., 2003) |

Following thermal cycling, positive and negative droplets were counted using the droplet reader (QX600™, Bio-Rad). Data was analyzed using Quantasoft analysis software (Bio-Rad), with automatic thresholding applied to distinguish positive and negative droplets across all samples.

Wells with <10,000 droplets were excluded from analysis. For gene expression analysis, RNA samples (previously diluted to 250-fold) were processed using One-Step RT-ddPCR Advanced kit for probes (Bio-Rad). Thermal cycling conditions were similar to DNA assay, with an additional reverse transcription step at 48 °C for 60 min followed by 40 cycles of PCR Table S4.

Table S4: Thermal cycling conditions for quantification of gene abundance and expression for ddPCR.

| **Steps** | **Temp. (°C)** | **Time** | **Cycling** |
|---|---|---|---|
| **Sample type: DNA; Target AOB & NOB** | | | |
| Enzyme activation | 95 | 10 min | 1 |
| Denaturation | 94 | 30 s | 39x |
| Annealing/Extension | 58.1 (AOB), 62.7 (NOB) | 2 min | |
| Enzyme deactivation | 98 | 10 min | 1 |
| **Sample type: RNA; Target AOB** | | | |
| Reverse transcription | 48 | 60 min | 1 |
| Enzyme activation | 95 | 10 min | 1 |
| Denaturation | 95 | 30 s | 39x |
| Annealing/Extension | 58.1 | 2 miin | |
| Enzyme deactivation | 98 | 10 min | 1 |
| Hold | 4 | | 1 |

#### 1.5.qPCR reaction mixture and thermal cycling condition

A 20 µL reaction volume contains Phusion Flash master mix (10 µL), primer mix (0.2 µL), nuclease-free water (4.8 µL), followed by DNA template (5 µL). PCR was performed as per the conditions indicated in Table S5. The amplicons along with the 100 bp ladder were run on an agarose gel (1.5%) using gel electrophoresis (Fisherbrand) immersed in TAE buffer for 20 mints at 100 volts. The products were visualized on agarose gel illuminator, the bands were excised and purified using gel purification kit (Purelink, Quick gel extraction kit K2100-12). The concentrations of the amplified DNA were determined with the Qubit using dsDNA high sensitivity assay (Invitrogen).

Table S5: Thermal cycling conditions for quantification of rpoB and 16S rRNA via qPCR.

| Steps | Temp. (°C) | Time | Cycling |
|---|---|---|---|
| **Thermal cycling conditions** | | | |
| Initial denaturation | 98 | 10 s | |
| Denaturation | 98 | 1 s | 30x |
| Annealing | 56.1 (rpoB), 55 (16S rRNA) | 5 s | |
| Extension | 72 | 7 s | |
| Final extension | 72 | 60 s | |
| **Thermal cycle for qPCR** | | | |
| Initial denaturation | 95 | 2 min | 1 |
| Denaturation | 95 | 5 s | 1 |
| Annealing | 56.1 (rpoB), 55 (16S rRNA) | 15 s | 38x |
| Extension | 72 | 15 s | |
| Final hold | 4 | | |
| Melt curve | | | |

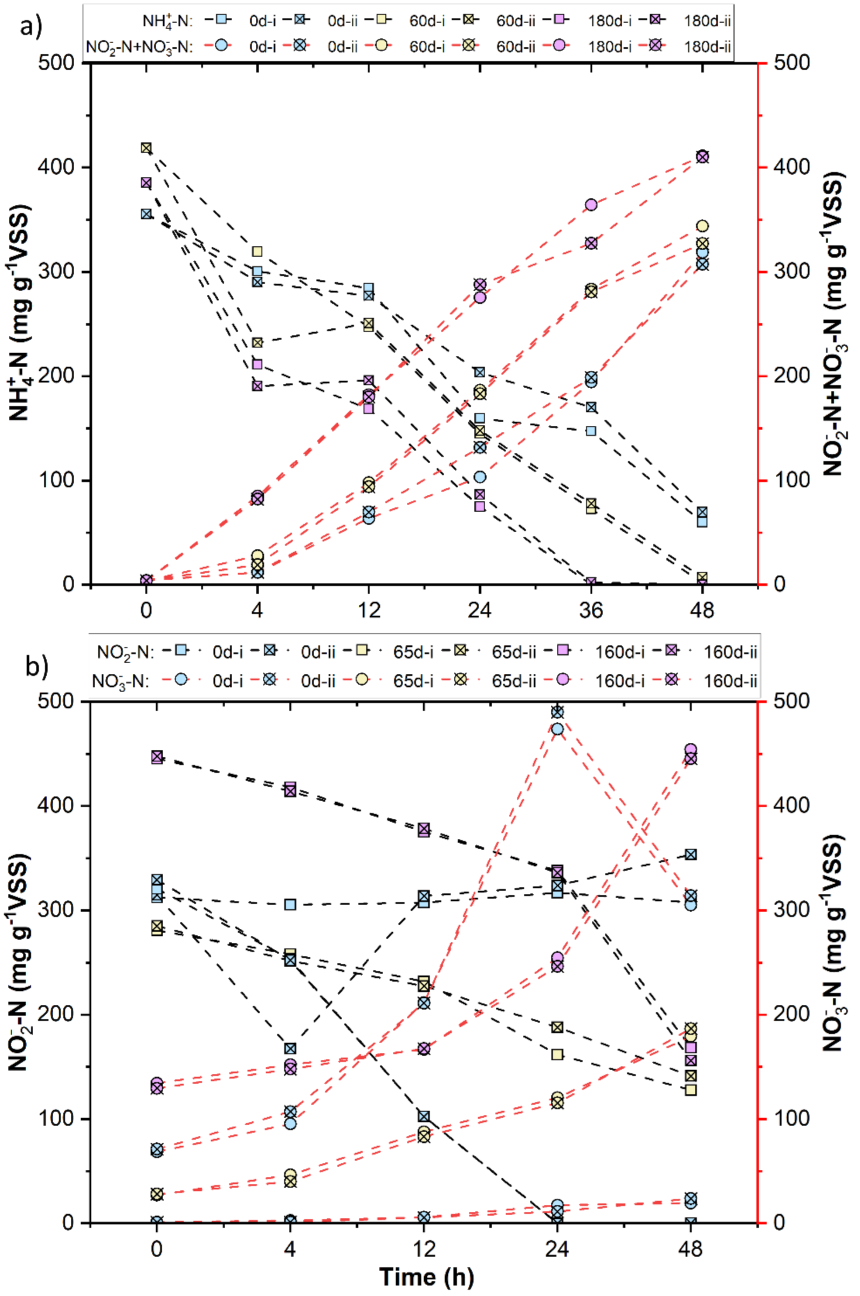


Figure S1. Batch test showing nitrogen transformation in (a) AOB-bioreactor, (b) NOB-bioreactor at various time points.

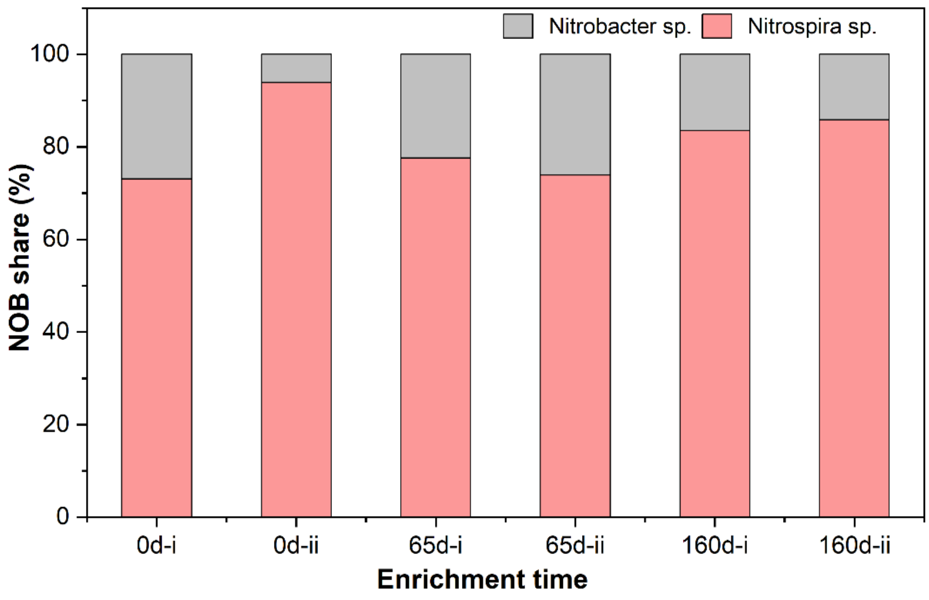


Fig. S2. Distribution of dominant NOB species across enrichment stages.

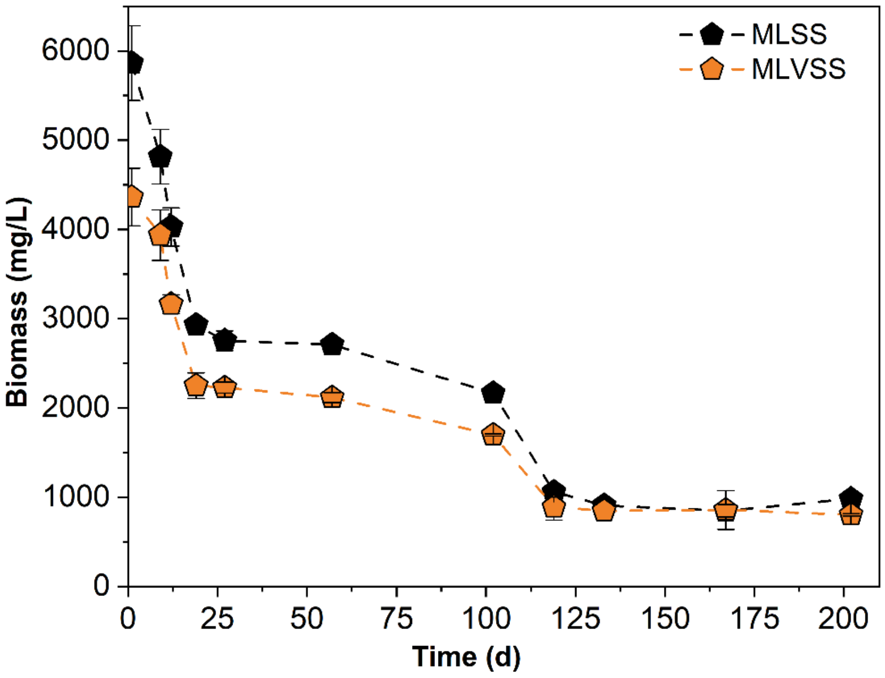


Figure S3. Biomass profiling in AOB-bioreactors.

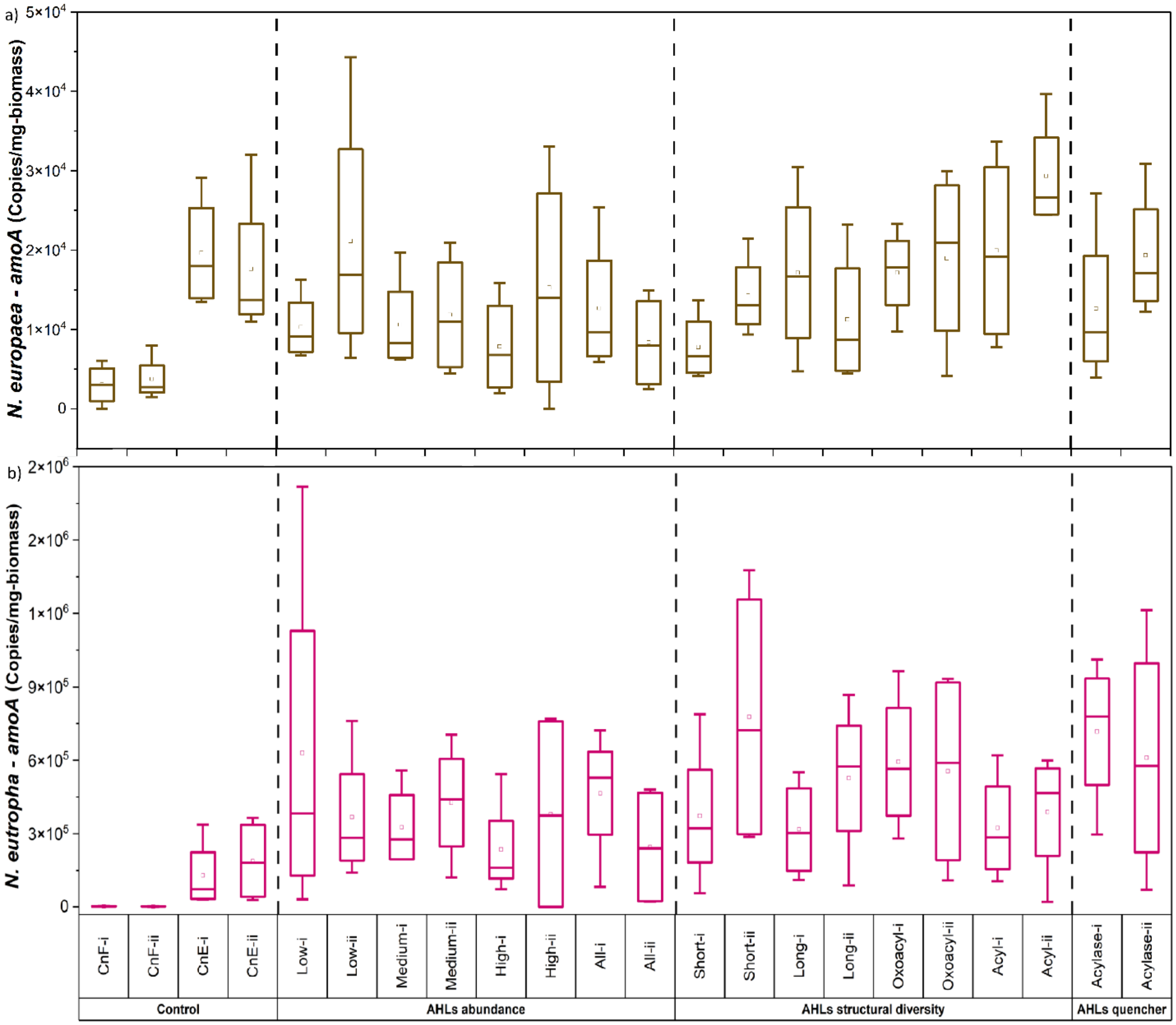


Fig. S4: AHLs influence across treatments with biological replicates indicating amoA gene abundance of (a) *N. europaea*, (b) *N. eutropha*.

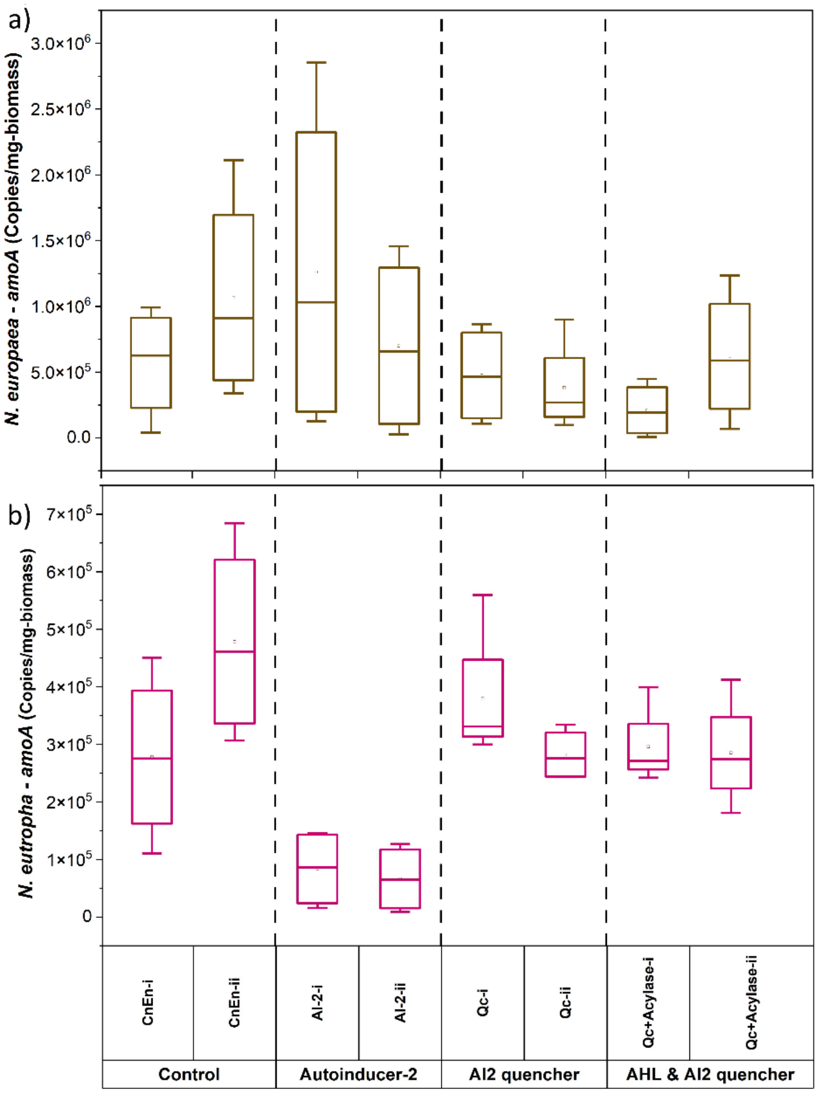


Fig. S5. AI-2 and quenching role across treatments with biological replicates indicating amoA gene abundance of (a) *N. europaea*, (b) *N. eutropha*.
